# The evolution of robustness and fragility during long-term bacterial adaptation

**DOI:** 10.1101/2025.01.24.632760

**Authors:** Doha Chihoub, Coralie Pintard, Richard E. Lenski, Olivier Tenaillon, Alejandro Couce

## Abstract

Most mutations affecting fitness are harmful, and their inevitable occurrence reduces mean population fitness. Theory predicts that well-adapted populations may evolve mechanisms to minimize this deleterious load. Direct selection to increase mutational robustness can be achieved in the laboratory. However, its spontaneous evolution during general adaptation remains uncertain, with mixed evidence across model systems. Here, we studied the effects of highly pleiotropic point mutations in *Escherichia coli* over a 15,000-generation adaptive trajectory. The fitness effects of both beneficial and deleterious mutations were attenuated with increased adaptation over time. In contrast, pleiotropic effects in new environments became more severe and widespread with greater adaptation. These results show that trade-offs between robustness and fragility can rapidly evolve in regulatory networks, regardless of whether driven by adaptive or non-adaptive processes. More broadly, these results show that adaptation can generate a hidden potential for phenotypic diversity, unpredictably shaping evolutionary prospects in new environments.

## INTRODUCTION

Populations may evolve mechanisms to improve the way they explore genotypic space. For example, in the early stages of adaptation to new environments, experimental populations often evolve high mutation rates (1) and substitute mutations with favorable epistatic profiles (2). Eventually, however, selection should favor mechanisms that counteract the influx of deleterious mutations, such as reducing mutation rates or increasing mutational robustness (i.e., buffering the effects of new mutations) (3). Reductions in mutation rates have been observed in well-adapted mutator populations (1), but evidence for the evolution of mutational robustness remains elusive.

It is certainly possible to directly select for variants with increased robustness by imposing strong mutagenesis (4), genetic drift (5), or thermal stress (6). However, it is unclear whether robustness can evolve spontaneously under typical circumstances, because its benefits are expected to be small and manifest only in well-adapted populations. Evolution experiments addressing this question are scarce and show mixed results: robustness has been observed to increase in viruses (7); decrease in viruses (8) and yeast (9); and show no directional change in bacteria (10) and yeast (11). These discrepancies may stem from differences in the starting genotype’s adaptation level, the type of mutations analyzed, or idiosyncrasies of the traits most relevant to fitness in each system.

An unexplored issue is whether robustness evolves to include the pleiotropic effects of new mutations. If so, does it apply only to those traits relevant to adaptation in the present selective environment, or does it extend to traits relevant anywhere? Examining pleiotropy could increase our ability to detect trends and help resolve conflicting empirical findings. To a first approximation, selection is expected to favor reduced pleiotropy and thereby confine the deleterious effects of new mutations to as few traits as possible – effectively mitigating their overall impact. For example, selection for robustness is likely stronger when pleiotropic effects alter traits coherently (effects on all traits are deleterious), rather than randomly (12).

Examining traits that have not been directly selected has a further advantage, insofar as it allows one to test the robustness-fragility trade-off hypothesis. That hypothesis suggests that resource allocation trade-offs that make systems robust to expected perturbations make them vulnerable to unexpected perturbations—a principle observed in engineering and proposed to apply to biological (13) and socioeconomic (14) systems.

## RESULTS AND DISCUSSION

To investigate changes in mutational effects along an established gradient of adaptation, we turned to the well-known long-term evolution experiment (LTEE) with *Escherichia coli* (15). We focused on three time points with evenly spaced fitness differences: the ancestor (“Anc”) and clones from 2,000 (“2K”) and 15,000 (“15K”) generations of the Ara-1 population, corresponding to fitness increases of ∼25% and ∼50%, respectively (Figure 1A) (15). We used site-directed recombineering to introduce the same set of 10 point mutations into the three genetic backgrounds (Figure 1B, Methods). We chose rifampicin-resistance mutations in the RNA polymerase (encoded by the *rpoB* gene), known for their wide-ranging fitness and pleiotropic effects (16).

**Figure 1.**
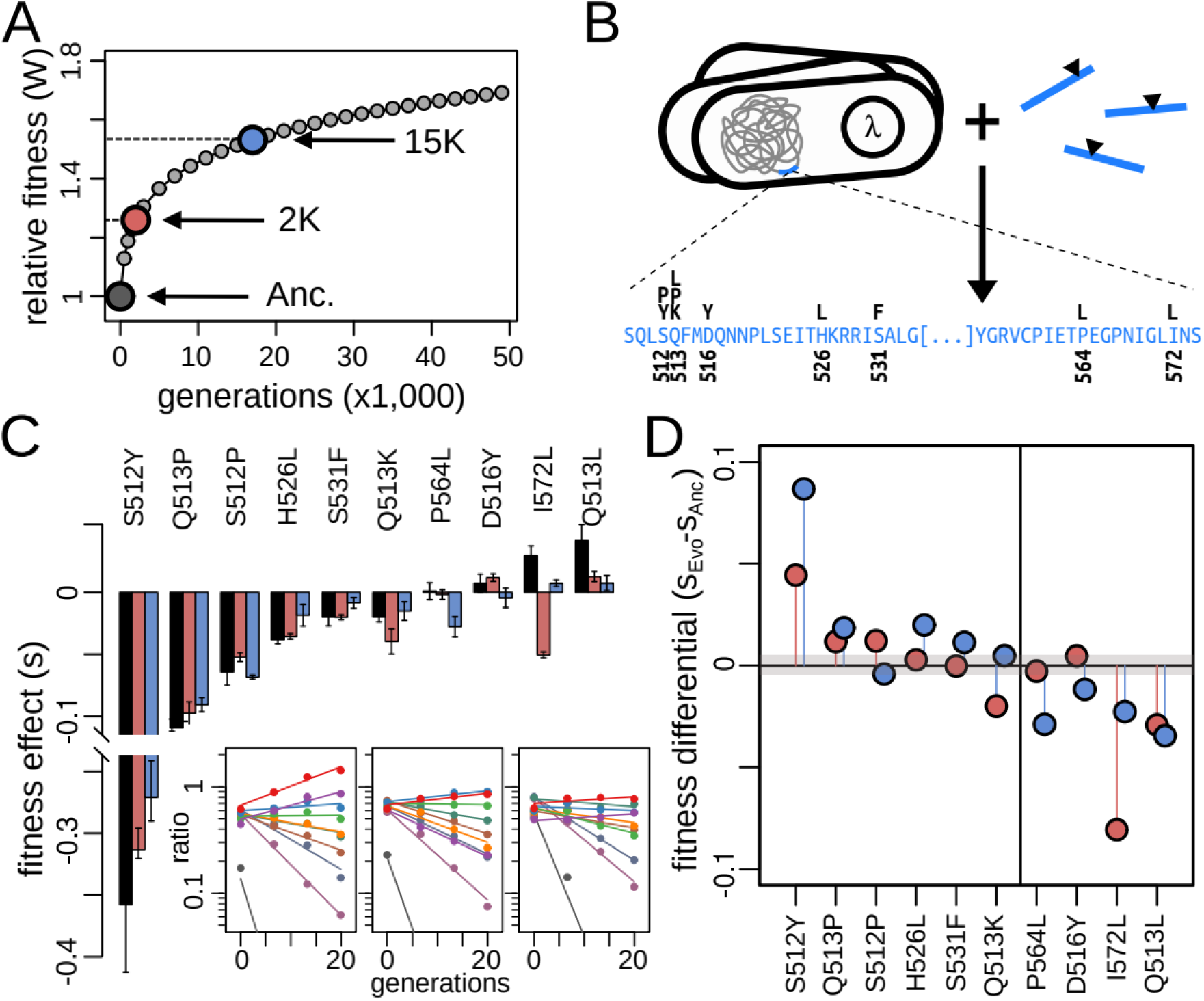
Fitness effects of RNA polymerase mutations in the LTEE environment. **(A)** Average fitness increase over time in the LTEE. We examined fitness effects in the ancestor and two evolved clones with fitness gains of ∼25% (2K) and ∼50% (15K). **(B)** Using lambda-red recombineering, we introduced the same 10 *rpoB* mutations, clustered within a 171-bp region, into the three genetic backgrounds. **(C)** Insets (left to right) show trajectories of mutant-to-wildtype ratios during bulk competitions for the Anc, 2K, and 15K backgrounds. Bar plot shows the fitness effects with standard errors estimated from those trajectories (colors: Anc, black; 2K, red; 15K, blue). **(D)** Difference in fitness effects in evolved strains (2K, red; 15K, blue) relative to the ancestor. Both deleterious and beneficial effects are progressively mitigated in the more-fit strains. Thin gray rectangle shows 95% confidence intervals from two replicate bulk competition experiments. Vertical line marks the approximate transition from deleterious to beneficial effects in the ancestral background.

Leveraging the close proximity of the 10 mutations (Figure 1B), we developed a targeted, deep-sequencing method to estimate fitness from allele frequency trajectories in bulk competition experiments (Methods). We pooled all mutants with their respective background strain (Anc, 2K or 15K) and tracked their trajectories during 3-day competitions under LTEE-like conditions (Figure 1C, inset; Methods). Both the magnitude and rank order of fitness effects were highly correlated across the three backgrounds (Figure 1C; Pearson’s r > 0.97; Spearman’s ρ > 0.78; P < 0.012, both cases). Among the 20 pairwise comparisons with the ancestor, we detected only three notable deviations, all involving a change in the sign of the fitness effects: I572L in the 2K background, and P564L and D516Y in the 15K background.

Despite the strong overall correlation, Figure 1C shows a tendency for effects to attenuate in the evolved backgrounds, particularly at the extremes (S512Y and Q513L). A plot of fitness differences between the evolved backgrounds and the ancestor clearly illustrates this trend, with both deleterious and beneficial effects becoming less so with adaptation (Figure 1D). We used linear regression to evaluate deviations from a one-to-one correspondence (i.e., slope β = 1). Slopes become progressively flatter in the evolved backgrounds (β = 0.8 at 2K; β = 0.72 at 15K; one-sided t-test, P < 0.01, both cases), supporting the evolution of mutational robustness along this adaptive trajectory.

We next investigated whether the observed attenuation was specific to the LTEE’s selective environment or extended to different conditions. To this end, we changed the culture medium from Davis Minimal Broth (DM) to Lysogeny Broth (LB), thus shifting the nutrient environment from a simple glucose-limited medium to a complex medium dominated by peptides along with small amounts of free amino acids, vitamins, and carbohydrates (17).

Mutation effects in LB were only weakly correlated, at best, with those in DM (Figure 2A; Pearson’s r = 0.29; Spearman’s ρ = 0.42; P > 0.2, both cases). Across the three backgrounds, however, the magnitude and rank order of effects were substantially conserved (Figure 2B; Pearson’s r > 0.67; Spearman’s ρ > 0 .71; P < 0.033, both tests, all three pairwise comparisons). In contrast to DM, fitness effects in LB showed no attenuation with adaptation during the LTEE (Figure 2C); the regression slopes are indistinguishable from a one-to-one correspondence (two-sided t-tests, P > 0.15). One explanation for this result is that the traits critical to fitness differ between DM and LB, with no trade-offs that induce fragility. Alternatively, partial overlap of critical traits, combined with trade-offs involving non-overlapping ones, could explain the absence of a directional trend. Directly investigating these possibilities is impractical, because the physiological traits most relevant for fitness in these media are unknown. A potentially illuminating approach is to study traits that can reasonably be assumed to be irrelevant under the LTEE conditions.

**Figure 2.**
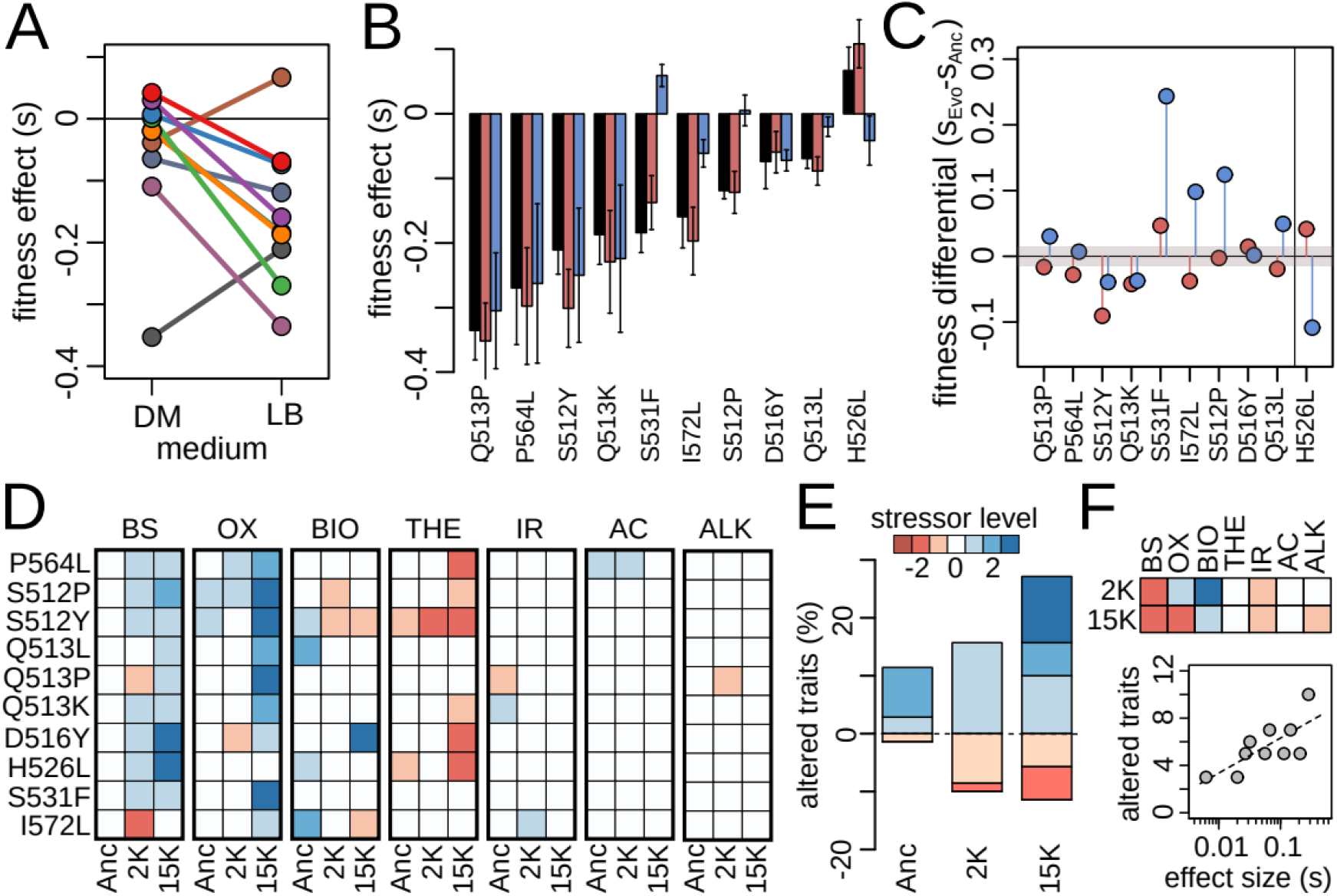
Pleiotropic effects of RNA polymerase mutations. **(A)** Mutations tend to be more deleterious, and their ranked effects change in LB compared to DM. **(B)** Fitness effects in LB are highly correlated across the three backgrounds, with a few conspicuous exceptions. Symbols as in Fig. 1C. **(C)** Fitness effects in better-fit strains show no clear directional change relative to the ancestor. Symbols as in Fig. 1D. **(D)** Heatmaps show changes in virulence-related traits caused by the 10 mutations across the three genetic backgrounds, recorded as changes in performance (red, decreases; blue, increases; scale in E shows the number of stressor levels by which mutations shift the point of growth inhibition, see SI). Traits include biofilm formation (BIO) and resistance to bile salts (BS), oxidative (OX), thermal (THE), iron starvation (IR), acidic (AC), and alkaline (ALK) stresses. **(E)** Trait alterations increase progressively as strains adapted to the LTEE environment, indicating a robustness-fragility trade-off. **(F)** Susceptibility profiles of the evolved backgrounds (heatmap). At the level of individual mutations, fragility correlates strongly with each mutation’s median fitness effect across backgrounds and media.

We thus examined several commonly accepted virulence-related traits (Methods, Figure 2D). We observed that mutations altered more of these traits in evolved backgrounds than in the ancestor (Figure 2E). This trend was marginally non-significant in 2K (9/70 vs. 18/70; chi-squared test, P = 0.087) but was pronounced in 15K (9/70 vs. 27/70; P = 0.001). This trend is partly driven by 15K’s increased susceptibility to bile salts and oxidative stress (Figure 2F), the most altered traits (Figure 2D). Finally, we observed that the 10 *rpoB* mutations vary in their tendency to exhibit pleiotropic effects (Figure 2D), which is well explained by their median absolute fitness impact across the three backgrounds and two media (Figure 2E; Pearson’s r = 0.73, P = 0.016).

Our results demonstrate that evolving populations can readily become robust to mutations affecting global regulatory networks. This phenomenon is likely missed by Tn-seq analyses because transposon insertions are more generally disruptive to cellular processes, perhaps explaining why a previous study of LTEE lines found no directional trend in robustness (9). We also report the evolution of fragility in traits that are probably unimportant for fitness in the LTEE conditions, demonstrating how adaptation can generate hidden phenotypic diversity with potential importance in alternative environments. While the emergence of this robustness-fragility trade-off matches predictions from systems control theory (13), we also note that the early stages of adaptation to new environments often involves suboptimal rewiring of global regulatory networks (18). Consequently, it remains to be determined whether these trade-offs are adaptive (i.e., resulting from selection to reduce deleterious load) or simply reflect the instability of a substantially altered transcriptome.

## MATERIALS AND METHODS

Site-directed mutagenesis in the *rpoB* gene was conducted using lambda-red-assisted recombineering. Targeted deep-sequencing was conducted on an Illumina MiSeq platform according to manufacturer’s specifications. Detailed materials and methods are provided in Supporting Information (SI).

## DATA, MATERIALS, AND SOFTWARE AVAILABILITY

All relevant data are provided in Dataset S1. Raw sequencing reads are deposited in the NCBI Sequence Read Archive (TBD).

## AUTHOR CONTRIBUTIONS

A.C. and O.T. conceived research; A.C., D.C. and C.P. performed research; R.E.L. contributed new reagents/analytic tools; A.C. analyzed data and prepared figures; A.C. drafted manuscript; A.C., R.E.L. and O.T. edited and critically revised the manuscript.

## ACKNOWLEDGMENTS

D.C. acknowledges support from the Algerian Ministry of Higher Education and Scientific Research under scholarship 029/Bis/PG/Espagne/2020-2021. R.E.L. acknowledges support from the U.S. National Science Foundation (DEB-1951307). O.T. acknowledges support from the *Agence Nationale pour la Recherche* (ANR-18-CE35-0005-0). A.C. acknowledges support from the *Agencia Estatal de Investigación* (Proyectos de I+D+i, PID2022-142857NB-I00; *Centros de Excelencia* “Severo Ochoa”, CEX2020-000999-S), and a *Comunidad de Madrid* “Talento” Fellowship (2019-T1/BIO-12882, 2023-5A/BIO-28940). We thank the Department of Genetics at Bichat-Claude Bernard Hospital (Paris) for access to their Illumina MiSeq platform, and Marion Réocreux for technical assistance with the system.

