## Supplemental information for "The evolution of robustness and fragility during long-term bacterial adaptation"

**This PDF file includes:**

SI Materials and Methods

Table S1

Supplementary References

### SI MATERIALS AND METHODS

#### Strains and media

We used the ancestral strain from the LTEE, *E. coli* B strain REL606 (1), and two derivatives isolated from the Ara-1 population at 2,000 (REL1164A) and 15,000 generations (REL7177A) (2). All three strains are unable to grow on arabinose as a carbon source ($Ara^-$). Bulk competition experiments were performed in Davis-Mingioli minimal medium supplemented with 25 µg/mL glucose (DM25, the standard medium in the LTEE) or Lysogeny Broth (LB; Miller’s modification). All other phenotypic assays were performed in LB. Selection for reversions to arabinose utilization ($Ara^+$) were conducted on M9 minimal salts agar plates supplemented with 1 mM L-arabinose.

#### Construction of *rpoB* mutants

Rifampicin-resistance point mutations known to affect fitness and influence multiple unrelated traits were introduced singly into REL606, REL1164A, and REL7177A using a double recombineering approach, as previously described (3). Briefly, we used the lambda-red recombinase system carried on plasmid pJk611, a pKD46 derivative containing a *sacB* gene for counter-selection on sucrose. This plasmid was first electroporated into electrocompetent recipient strains, and transformants were selected on LB agar plates containing 100 µg/mL ampicillin; the plasmid backbone carries an ampicillin-resistance cassette. Individual colonies were then isolated and stored at −80°C. Next, pJk611-carrying strains were made electrocompetent again and electroporated with two 70-bp oligonucleotides per rifampicin-resistance mutant (Table S1). The first oligonucleotide is designed to introduce the desired point mutation in *rpoB*, while the second introduces a A275G mutation in *araA*. This mutation reverts a loss-of-function mutation present in REL606 (1) and its evolved derivatives, restoring the ability to grow on arabinose as the sole carbon source ($Ara^+$). Selection for this reversion enriches for cells that are competent, express the recombineering genes, and incorporate the oligonucleotides—thereby increasing the efficiency of site-directed mutagenesis at the desired *rpoB* site. To screen for these revertants, cells were plated on M9 agar supplemented with L-arabinose and incubated for 48 h. Three isolated revertants per mutant were re-streaked onto LB plates with 100 µg/mL rifampicin, and $Ara^+$ $Rif^r$ colonies were subjected to colony PCR and Sanger sequencing to confirm the presence of the desired *rpoB* mutation (Table S1).

#### Bulk competition experiments

Overnight cultures of all 10 mutants were pooled with their respective background strain (Anc, 2K, or 15K) and propagated for 3 days under conditions mimicking the LTEE: daily 1/100 dilutions in 50-mL flasks containing 10 mL of medium, incubated at 37°C in an orbital shaker (1). For competitions in DM25, a 100-µL aliquot was taken each day to seed the next flask; the remaining culture was pelleted using a Beckman LS60 ultracentrifuge (30 min, 48,000×g) and subjected to genomic DNA extraction with the DNeasy Blood & Tissue Kit (Qiagen). For competitions in LB, 1 mL of culture was used for each genomic DNA extraction. Daily counts of viable cells confirmed that each passage conformed to the expected ~6.6 generations of binary fission given the 100-fold dilutions and regrowth.

### Targeted deep-sequencing

Genomic DNA from each sample was homogenized after quantification using a Qubit dsDNA HS Kit (Thermo Fisher). A 260-bp region of *rpoB*, containing the 10 sites where mutations were introduced, was then amplified by PCR using the 59- and 60-bp primers listed in Table S1. Each primer included a 20-bp sequence homologous to the flanks of the *rpoB* target region, a 6-bp degenerate region to aid clustering on Illumina sequencing platforms, and a 33- or 34-bp sequence compatible with the Nextera XT DNA Library Prep Kit (Illumina). After band purification using the QIAquick Gel Extraction Kit (Qiagen), we ran a second PCR to add indexes for multiplexed, paired-end sequencing. This secondary PCR involved an 18-cycle PCR using the Nextera XT primers and buffers. The final libraries were purified, quantified, and sent for sequencing on a MiSeq platform (Bichat Hospital, Paris, France).

### Fitness estimation

To obtain high-resolution fitness estimates from the targeted sequencing reads, we first used a Python script to compare paired-end reads and generate consensus sequences. Discrepancies between forward and reverse reads were conservatively resolved by retaining only the wild-type sequence. A second Python script scanned the consensus reads to count perfect matches to any of the 10 possible mutations or the wild-type sequence. The $\sim 1.5 \times 10^5$ raw reads per sample from MiSeq sequencing were reduced to $\sim 0.5 \times 10^5$ reads after processing with these two stringent scripts. Next, mutant counts were processed in R to estimate fitness as the slope of mutant frequency trajectories relative to the wild-type trajectory. This slope, expressed as the natural logarithm, approximates the selection coefficient ($s$) from classical population genetics and is related to the ratio of realized growth rates ($W$), the standard fitness metric in the LTEE literature, by a scaling factor of ln(2) (4). Slopes were estimated using linear regression implemented with the *lm()* function in R, incorporating weights proportional to read counts at each time point to reduce noise from low-count observations (5). Initial read counts averaged 46,220 ± 17,488 (mean ± standard deviation) across all mutant and wild-type alleles in the three backgrounds and two growth media. Despite declining numbers of some mutants during the competitions (up to a maximum of ~500-fold), the read counts consistently remained above 30 per sample.

### Phenotypic assays

All assays were performed by adding $\sim 10^4$ colony-forming units from overnight cultures to 96-well microtiter plates filled with 100 μL of LB (supplemented as required).

Growth was quantified after incubation using a SPECTROstar Nano (BMG Labtech, Germany). Depending on the assay, plates were incubated for 24 or 48 h and optical density (OD) readings were performed at 550 nm or 600 nm. Each experiment included at least five biological replicates. For all phenotypes except biofilm formation, susceptibility was determined by exposing cultures to a range of stressor levels and recording the maximum level at which growth was detectable (based on OD exceeding three times the standard deviation of blank wells). Alteration scores were expressed as the number of stressor levels by which mutants differed from the ancestral background. Oxidative stress was induced with paraquat at doubling concentrations from 8 to 64 µg/mL. Bile salt tolerance was tested at concentrations ranging from 0.63% to 8%. Iron starvation was induced using bipyridyl at doubling concentrations from 0.125 to 2 mM, with experimental vessels thoroughly washed to remove trace iron. Acid and alkaline stresses were imposed by adjusting LB to pH levels of 3, 3.5, 4, 4.5, 5 (acidic) and 8, 8.5, 9, 9.5, 10 (alkaline). Temperature susceptibility was assessed at seven temperatures from 40°C to 46°C, incremented by 1°C. For biofilm formation, cultures were incubated for 48 h at 28°C, washed three times with distilled water, stained with crystal violet, dissolved in acetic acid/ethanol (20:80), and subjected to OD measurements at 550 nm (6). Alteration scores were categorized as 25%, 50%, or 75% deviations (in either direction) in biofilm formation compared with the ancestral background.

**Statistical analyses and data visualization**

All statistical analyses were performed with the R programming language (version 3.6.3), using built-in functions and available packages (7). Correlation analyses, linear regressions, t-tests, and logistic regressions were performed with the functions *cor.test()*, *lm()*, *t.test()*, *qt()*, and *glm()*, respectively. Data visualization for Figure 2D was performed using the 'pheatmap' library (8), while standard base R plotting functions were used elsewhere.

**Table S1. Primers used in this study**

| Name | Sequence[1] |
|---|---|
| araA_D92G | 5'-CACACCTTCTCCCCGGCCAAAATGTGGATCAACGgCCTGACCATGCTCAACAAACCGTTGCTGCAATTCC-3' |
| rpoB_S512P | 5'-CTCAGACAGCGGGTTGTTCTGGTCCATAAACTGAGgCAGCTGGCTGGAACCGAAGAACTCTTTCACTGCT-3' |
| rpoB_S512Y | 5'-TCTCAGACAGCGGGTTGTTCTGGTCCATAAACTGAtACAGCTGGCTGGAACCGAAGAACTCTTTCACTGC-3' |
| rpoB_Q513L | 5'-TAATCTCAGACAGCGGGTTGTTCTGGTCCATAAACaGAGACAGCTGGCTGGAACCGAAGAACTCTTTCAC-3' |
| rpoB_Q513P | 5'-TAATCTCAGACAGCGGGTTGTTCTGGTCCATAAACgGAGACAGCTGGCTGGAACCGAAGAACTCTTTCAC-3 |
| rpoB_Q513K | 5'-AATCTCAGACAGCGGGTTGTTCTGGTCCATAAACTtAGACAGCTGGCTGGAACCGAAGAACTCTTTCACT-3 |
| rpoB_D516Y | 5'-TTTGTGCGTAATCTCAGACAGCGGGTTGTTCTGGTaCATAAACTGAGACAGCTGGCTGGAACCGAAGAAC-3' |
| rpoB_H526L | 5'-GACCGCCTGGGCCGAGTGCGGAGATACGACGTTTGaGCGTAATCTCAGACAGCGGGTTGTTCTGGTCCAT-3' |
| rpoB_S531F | 5'-CACGTTCACGGGTCAGACCGCCTGGGCCGAGTGCGaAGATACGACGTTTGTGCGTAATCTCAGACAGCGG-3 |
| rpoB_P564L | 5'-ACAGAGAGTTGATCAGACCGATGTTCGGACCTTCAaGGGTTTCGATTGGACATACGCGACCGTAGTGAGT-3 |
| rpoB_I572L | 5'-TTCGTTAGTCTGTGCGTACACGGACAGAGAGTTGAgCAGACCGATGTTCGGACCTTCAGGGGTTTCGATT-3 |
| rpoB_IllumF | 5'-TCGTCGGCAGCGTCAGATGTGTATAAGAGACAGNNNNNNAGCAGTGAAAGAGTTCTTCG-3 |
| rpoB_IllumR | 5'-GTCTCGTGGGCTCGGAGATGTGTATAAGAGACAGNNNNNNGGAAGCCGTATTCGTTAGTC-3 |
| rpoB_SangF | 5'-AGCAGTGAAAGAGTTCTTCG-3 |
| rpoB_SangR | 5'-GGAAGCCGTATTCGTTAGTC-3 |

[1]In the site-directed recombineering primers (one for *araA* and 10 for *rpoB*), the nucleotide that introduces the mutation is shown in lowercase.
